# Multiple and Dissociable Effects of Sensory History on Working-Memory Performance

**DOI:** 10.1101/2021.10.31.466639

**Authors:** Jasper E. Hajonides, Freek van Ede, Mark G. Stokes, Anna C. Nobre, Nicholas E. Myers

## Abstract

Behavioural reports of sensory information are biased by stimulus history. The nature and direction of such serial-dependence biases can differ between experimental settings – both attractive and repulsive biases towards previous stimuli have been observed. How and when these biases arise in the human brain remains largely unexplored. They could occur either via a change in sensory processing itself and/or during post-perceptual processes such as maintenance or decision-making. To address this, we analysed behavioural and magnetoencephalographic data from a working-memory task in which participants were sequentially presented with two randomly oriented gratings, one of which was cued for recall at the end of the trial. Behavioural responses showed evidence for two distinct biases: 1) a within-trial repulsive bias away from the previously encoded orientation on the same trial, and 2) a between-trial attractive bias towards the task-relevant orientation on the previous trial. Multivariate classification of stimulus orientation revealed that neural representations during stimulus encoding were biased away from the previous grating orientation, regardless of whether we considered the within- or between-trial prior orientation – despite opposite effects on behaviour. These results suggest that repulsive biases occur at the level of sensory processing and can be overridden at post-perceptual stages to result in attractive biases in behaviour.

## 1 Introduction

Stimulus history modulates performance guided by sensory input. Reliance on temporal correlations is deeply engrained in the visual system (Simoncelli & Olshausen, 2001) and can be beneficial to guide perception within a world that is largely stable over short time scales (Dong & Atick, 1995). Past sensory evidence can be an effective prior for extracting signals from the noisy sensory stream and to help maintain a stable representation to bridge blinks, eye movements, or visual occlusions.

Recent studies have also revealed robust short-term influences of prior stimulation in delayed-response and working-memory tasks (Bae & Luck, 2017; Cicchini, Anobile, & Burr, 2014; Cicchini, Mikellidou, & Burr, 2018; Czoschke, Fischer, Beitner, Kaiser, & Bledowski, 2019; Fischer & Whitney, 2014; Fritsche, Mostert, & de Lange, 2017; Huang & Sekuler, 2010). More specifically, remembered items are often reported as more *similar* to the task-relevant stimulus in a previous trial. This attractive bias, sometimes referred to as the serial-dependency bias, is suggested to increase temporal stability by acting as a prior (Cicchini, Anobile, & Burr, 2014; Fischer, Czoschke, et al., 2020; Fischer & Whitney, 2014; Kiyonaga, Scimeca, Bliss, & Whitney, 2017). Conversely, features of items sequentially presented within trials are often judged as more *dissimilar* to each other (Born & Tootell, 1992; Fritsche, Mostert, & de Lange, 2017; Störmer & Alvarez, 2014). This repulsive bias could result from efficient coding of temporally autocorrelated signals (Cicchini, Mikellidou, & Burr, 2018). The amplification of subtle differences between stimuli could additionally serve to optimise perceptual decision-making (Cicchini, Mikellidou, & Burr, 2018; Kiyonaga, Scimeca, Bliss, & Whitney, 2017; van Bergen & Jehee, 2019). Despite their opposite directions, attractive and repulsive performance biases have been shown to jointly influence task processing (Czoschke, Fischer, Beitner, Kaiser, & Bledowski, 2019; Fritsche, Mostert, & de Lange, 2017; Fritsche, Spaak, & de Lange, 2020; Sadil, Cowell, & Huber, 2021), albeit at different time scales (Gekas, McDermott, & Mamassian, 2019; Sheehan & Serences, 2021). While the current literature has largely focused on behaviour, relatively less work has examined whether these opposing biases share a neural mechanism, and in particular whether both biases may arise at early or late sensory processing stages. A number of studies have shown evidence for between-trial biases arising in early visual cortex (Sheehan & Serences, 2021; St. John-Saaltink, Kok, Lau, & De Lange, 2016), but with insufficient temporal resolution to isolate early processing stages, or they have measured activity only in frontal cortex but not sensory cortex (Papadimitriou, White, & Snyder, 2017).

The current study combined behavioural reports in a working-memory task and continuous magnetoencephalographic (MEG) recordings to assess within- and between-trial biases induced by sensory history with high temporal resolution. We devised a task in which participants were sequentially presented with two orientations and used a continuous, precision response to reproduce one of them at the end of a trial. Additionally, we included a cue that indicated which of the two orientations had to be reported in the trial to test for possible differential susceptibility of biases to the task-relevance of stimuli (as per Bae & Luck, 2020; Fischer, Czoschke, et al., 2020). We compared systematic biases induced by the first orientation on the second (within-trial bias) as well as biases from the previous trial on the current trial (between-trial bias). Critically, we tested whether we could find neural signatures for such biases using a decoding approach that leveraged the high temporal resolution of MEG. We set out to decode the presented orientation and expected to find the within- and between-trial biases in the neural data that mirrored the behavioural biases.

Previewing the results, we confirmed that behavioural responses were systematically pushed away from previous orientations on the same trial and pulled toward the orientations recalled on the previous trial. Mirroring the behavioural repulsion results, the sensory neural representation of stimulus orientation was shifted away from the preceding orientation presented on a given trial. However, we found no neural evidence for an attractive between-trial sensory bias during orientation encoding, despite such a bias observed in behaviour. Instead, we always observed a repulsive neural bias, regardless whether we considered orientations from the same or previous trial as a source of the bias.

## 2 Methods

### 2.1 Participants

Twenty healthy volunteers with normal or corrected-to-normal vision participated in the study. All participants were between 20 and 36 years old (mean 25.4 years old; eleven females). Prior to taking part in the study, volunteers provided their informed consent according to the procedures approved by the Central University Research Ethics Committee of the University of Oxford. Participants received £15 per hour compensation for taking part in this study.

### 2.2 Experimental set-up

Participants sat in the MEG scanner, which was situated in a dimly lit, sound-proof, and magnetically shielded room. A projection screen was placed at a viewing distance of 90 cm. Visual stimuli were projected at the back of the screen at a spatial resolution of 1024 × 768 pixels using a refresh rate of 60 Hz using a Panasonic DLP projector (PT-D7700E).

The task was programmed and presented using Matlab (Mathworks, Nantick, WA) in conjunction with the Psychophysics Toolbox (Brainard, 1997). Participants indicated their responses on an optic-fibre response box.

### 2.3 Task

Participants performed a precision working-memory task in which they reproduced the orientation of one of two grating stimuli presented sequentially with independent orientations (**Figure 1**). Simultaneously with the presentation of the second grating, participants were cued as to which grating orientation to report when probed at the end of the trial. In half of the trials, only the first or second grating was presented and used for reporting.

**Figure 1:**
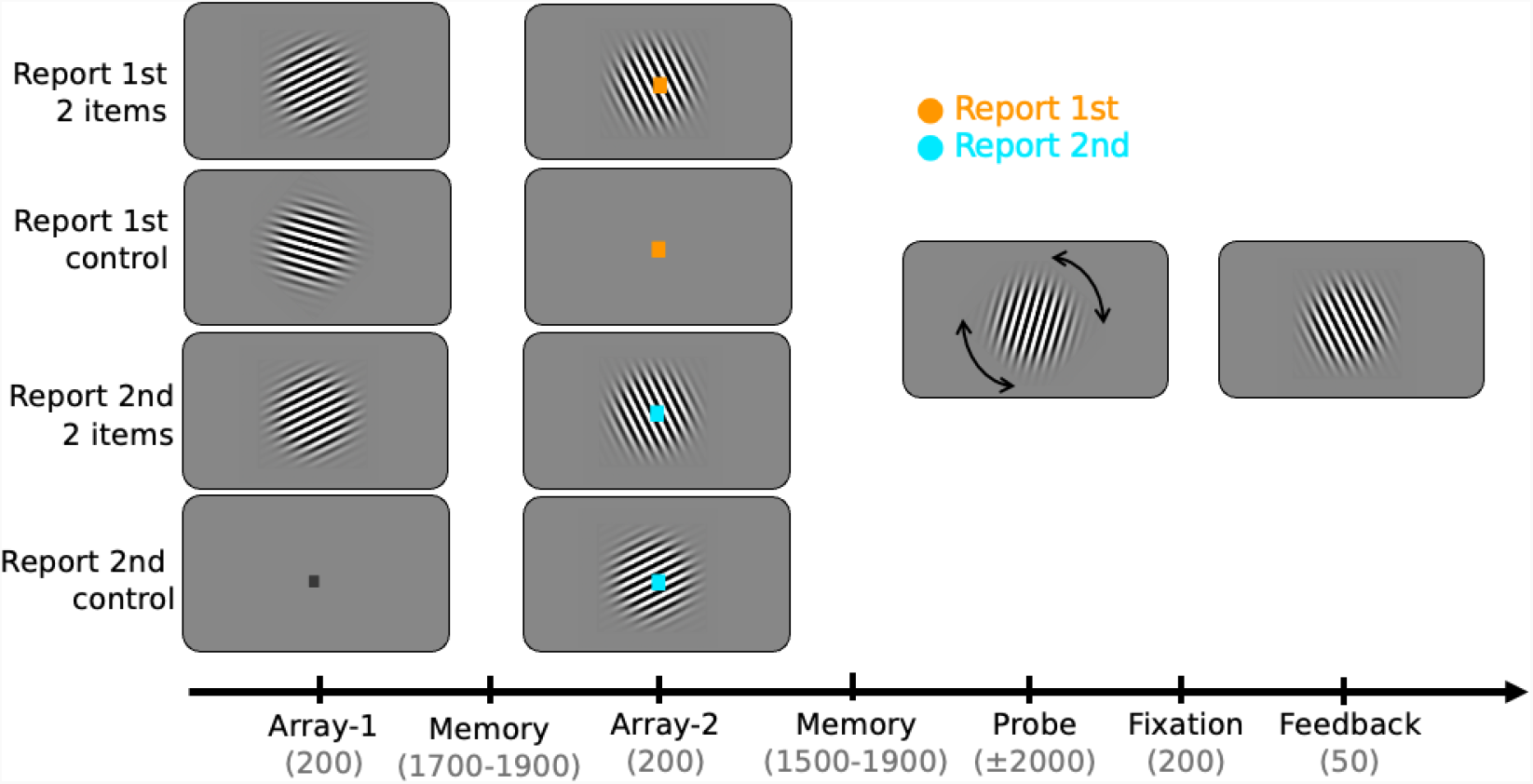
Experimental task design. Participants were presented with two arrays that contained an oriented bar grating or a place holder fixation dot. An oriented grating was presented either in the first (25% of trials), second (25%), or both (50%) arrays. A colour cue presented on array 2 indicated the relevant stimulus for recall. The cue size in array-2 is enlarged for visualisation purposes. The colour cued either the first array or the second array. On trials with only one orientation presented, the cue was redundant but still presented. Mappings between colour and cue meaning were counterbalanced. After matching the orientation in the probe display to the orientation in memory, participants were presented with feedback in the form of a grating with the correct orientation. Timings in parentheses are in milliseconds.

Each trial started with a central fixation dot (0.2° visual angle) on screen for 800 ms with a grey background (RGB: 127, 127, 127; **Figure 1**). Subsequently, a sinusoidal Gabor stimulus at a random angle was centrally presented on the visual display for 200 ms (diameter of 6° visual angle,2 cycles per degree of visual angle, 50% contrast, tapered by a Gaussian envelope with a 1.5° standard deviation). On trials in which the first grating was not presented the fixation dot changed colour from black to grey (RGB: 192, 192, 192) to signal the omission. After a delay of 1700 – 1900 ms, the second grating was presented. This second grating had the same properties as the first grating except for the angle, which was randomly drawn independently from that of the first grating orientation. At the centre of the grating, the fixation dot changed colour (orange: (255, 161, 0); cyan: (0, 236, 255)). The colour signalled if the first orientation (report 1st trial) or the second orientation (report 2nd trial) would be probed. Cue-colour contingencies of the experiment were counterbalanced across participants and changed halfway through the experiment. After the colour contingencies switched, participants practised the new contingencies over the course of one block before continuing with the second half of the session. The colour cue was always valid in indicating which grating orientation was relevant for the reporting stage at the end of the trial. On trials in which the second grating was absent the cue would therefore always indicate the first orientation.

After another delay of 1700 – 1900 ms, a probe grating appeared. Participants had to adjust its orientation to match the cued grating in memory. Adjustments were made by pressing buttons with the right hand to rotate the grating either clockwise (middle finger) or anticlockwise (index finger). Responses were confirmed by pressing a button with the left index finger. A 200-ms fixation period followed, after which participants were presented with 50 ms of feedback in the form of a grating indicating the correct orientation.

In total, participants completed 400 trials. In half of the trials, two items were presented in 200 trials (100 trials with report 1st cue and 100 with report 2nd cue). Only one item was presented in the remaining 200 trials (100 with only grating 1 and 100 only grating 2). The resulting factorial design (relevant grating, number of gratings presented) included four conditions: report 1st with two items presented, report 1st with one item presented, report 2nd with two items presented, and report 2nd with one item presented. Trial types were randomly mixed and presented in blocks of 50 trials, with each block lasting approximately 10 minutes.

### 2.4 Behavioural Analysis

Response error was quantified by calculating the circular distance between the recalled orientation and the cued orientation. All responses were mapped onto a -90° to 90° space to compute circular error. In all between-trial analyses, we excluded the first trial of each block.

### 2.5 Mixture Modelling

We fit a classical mixture model frequently used in the working memory literature to investigate the relative rate of responses to targets, guesses, and erroneous responses to the wrong target (‘swap errors’ Bays, Catalao, & Husain, 2009; Schneegans & Bays, 2017). The model was fit separately for each condition (report 1st in 2-item trials; report 1st in 1-item control; report 2nd in 2-item trials; report 2nd in 1-item control). The mixture model estimated the precision of the von Mises distribution, target response rate, guess rate, and swap rate to the item that was presented on the same trial but not cued (only 2-item trials). After fitting the data to the entire dataset, the model provided single-trial weights for target response rate, guess rate, and swap rate.

### 2.6 Performance Bias Calculation

We calculated the performance bias (the signed circular difference between the response and the target orientation) as a function of the circular difference between the target orientation and the orientation that induced the bias (the target orientation in the preceding trial or the task-irrelevant orientation in the same trial). We computed the difference between the target orientation and inducer through subtraction, with all angular differences mapped between -90° and 90°. These distances were binned into 64 equally sized, overlapping bins, with each bin containing 25% of trials. We computed the average signed error (bias) across all trials within each bin. Edge artefacts were avoided by wrapping the angular differences around, to ensure the average was computed over a constant number of trials and centred on the midpoint of the bin. Subsequently, we sign-flipped the performance bias in bins with a negative distance and averaged over negative (−90° to 0°) and positive distances (0° to 90°) between target orientation and inducing orientation, resulting in 32 bins. The summed bias over absolute distances (0° to 90°) calculated over the 32 bins for each condition and participant served as a measure of bias.

### 2.7 MEG Acquisition

Participants were seated in the MEG scanner after being instructed about the task specifics. They completed one practice block while seated in the scanner prior to MEG recording onset. Participants were instructed to maintain their gaze at the central fixation dot and to minimise blinking throughout the trial.

Neuromagnetic data were acquired using a whole-head VectorView system including 204 planar gradiometers and 102 magnetometers (Elekta Neuromag Oy, Helsinki, Finland) in a magnetically shielded room. Throughout the experiment, participants’ head position was monitored continuously using index coils placed at four points on the head. Magnetic field strength was sampled at a rate of 1000 Hz and band-pass filtered on-line between 0.03 Hz and 300 Hz. In addition, vertical and horizontal electro-oculograms were measured using electrodes placed above, below, and adjacent to the eyes. Eye movements were monitored using an EyeLink 1000 (SR Research) eye tracker at a frequency of 1000 Hz.

### 2.8 MEG Data Preprocessing

The data were pre-processed offline using Fieldtrip (Oostenveld, Fries, Maris, & Schoffelen, 2011), OHBA software library (OSL) drawing on SPM8 (http://www.fil.ion.ucl.ac.uk/spm), and Elekta software. Prior to any preprocessing, the MEG data were visually inspected to remove and interpolate any sensors that displayed excessive levels of noise and were subsequently de-noised and motion corrected using Maxfilter Signal Space Separation (Taulu, Kajola, & Simola, 2004) before removing independent components related to cardiac and eye-blink artefacts. Data were epoched around the first grating and second grating (from 400 ms prior to grating onset to 900 ms after onset) and downsampled to 200 Hz. Trials with high variance in either gradiometers or magnetometers were identified and excluded using a generalised ESD (extreme studentised deviate; Rosner, 1983) test at a 0.05 significance threshold. This resulted in 7.49% ± 11.55% (mean ± standard deviation) being excluded during preprocessing.

### 2.9 LDA Classification

Data were further pre-processed. Magnitudes of magnetometers were approximately matched to gradiometers by multiplication (factor 20) and subjected to spatio-temporal decoding as described in (Hajonides, Nobre, van Ede, & Stokes, 2021; Wolff, Jochim, Akyürek, Buschman, & Stokes, 2020; Wolff, Jochim, Akyürek, & Stokes, 2017). Data from all 306 MEG sensors across a sliding window of 30 time points (150 ms) were concatenated into a vector. Pre-stimulus baselining was not applied to maintain stable information from previously presented stimuli. Dimensionality was reduced for each time point through a principal component analysis, maintaining 90% of the variance (between 250 to 600 ms this was around 209 ± 39 components, mean ± std.). To train an LDA classifier, the data were split into training and testing sets using 10-fold stratified cross-validation. Grating angles were binned into 10 equally spaced orientation bins (0° to 18°, 18° to 36°, 36° degree to 126°, 126° to 144°, 144° to 162°, 162° to 180°). For each trial and time point, we thus obtained 10 LDA distances estimating the likelihood for each of the bins. Representational similarity curves were constructed by aligning evidence across trials around the same category bin. In cross-decoding analyses, LDA classifiers were trained on orientation bins of one event (e.g., presented grating) but classifier evidence aligned around bins of another orientation (e.g., target orientation on the previous trial). The resulting representational similarity curves were convolved with a cosine.

To test which sensors most significantly contributed to the classifier likelihoods observed in our multivariate methods, we also ran a searchlight decoding analysis (Kriegeskorte, Goebel, & Bandettini, 2006). In this analysis, we iteratively considered a small group of sensors and were thereby able to map the approximate locus of the observed effect. More specifically, we selected data from each sensor plus its 47 most closely adjacent neighbours (magnetometers and gradiometers included) and ran the same classification analysis as described above.

### 2.10 Calculation of the Neural Asymmetry Score as a measure of Neural Bias

For within-trial biases, we assessed processing of the second grating and only considered 2-item trials. The classifier was trained on all presentations of the second grating and bin likelihoods were generated for each trial. For between-trial analyses, we analysed orientation processing of both the first and second grating in the current trial. For this reason, we trained the classifier on all trials and generated bin predictions for all trials.

Subsequently, based on the results from the performance-bias analyses, we selected trials in which the angular distance between the inducer and the grating orientation on the display led to a significant behavioural bias at the group level. In the case of the within-trial repulsive bias, the inducer was the orientation of the first grating on the same trial. For the between-trial analyses, the inducer was the target orientation reported on the previous trial (except for control analyses, where the unreported orientation was used as the target). As a dependent variable, we considered likelihood estimations for each orientation bin, where we expect the highest likelihood for the angular bin that has zero offset to the presented orientation and decreasing likelihoods for bins with larger angular distances to the presented orientation. We separately assessed likelihood estimations for trials in which the inducer orientation was clockwise (CW) vs counterclockwise (CCW) with respect to the current orientation. For both CW and CCW trials, we separately averaged the evidence from the orientation bins CW (−72° to -18°) and evidence from the CCW bins (18° to 72°). Asymmetry scores were computed by obtaining the difference between the two groups of angular bins (CW minus CCW). Finally, we calculated an overall neural bias score by subtracting asymmetry scores on trials with CW vs. CCW inducers. Attractive neural biases resulted in a positive score (i.e., trials with CW angular distances resulted in more CW evidence, CCW angular distances resulted in more CCW evidence), whereas repulsive neural biases resulted in a negative score (i.e., CW angular distances resulted in less CW evidence than CCW trials, and vice versa).

### 2.11 Statistical Testing

Statistical tests were computed using both JASP (Team, 2020) and Scipy (Virtanen et al., 2020).

We tested the time series of cosine-convolved classifier evidence against zero using a cluster-based permutation test, which addresses the multiple comparison problem (using MNE; Gramfort et al., 2013). We ran 100,000 iterations. The clusters with groups of time points significantly different from zero are indicated in the relevant figures using horizontal lines. Cluster-based permutation testing was also applied to performance bias across angular distance between the presented orientation and the inducer orientation.

The time period of interest for the neural bias analyses included all timepoints in which cosine-convolved classifier evidence for the presented grating orientation was significantly above zero in our comparison analyses (in all reported time averages, time points between 250 and 600 ms were used).

All tests were two-sided unless stated otherwise.

## 3 Results

### 3.1 Error Rates

Participants were accurate in reproducing the target orientation (mean response error 11.73° ± 0.70° SEM; mean standard deviation 17.61° ± 1.07° S.E.M., see Table 1 for condition-wise performance). A 2-by-2 repeated-measures ANOVA on response error showed main effects of cue type (*F* _1,19_ = 16.49, *p <* .001, *η*^2^ = 0.374) and number of stimuli presented (*F* _1,19_ = 29.78, *p <* .001, *η*^2^ = 0.075). Cue type was significant for both 1- or 2-item trials, with absolute error higher on report 1st than on report 2nd trials for both 2-item trials (*t*_19_ = 3.972, *p <* .001, *d* = 0.888; see **Table 1**) and 1-item trials (*t*_19_ = 3.948, Bonferroni-corrected *p <* .001, *d* = 0.883). By contrast, number of items presented primarily affected the report 1st conditions. Error was higher on report 1st 2-item than on report 1st 1-item trials (*t*_19_ = 5.665, *p <* .001, *d* = 1.267) but did not significantly differ between report 2nd 2-item and report 2nd 1-item trials (*t*_19_ = 1.885, *p* = .075, *d* = 0.421), leading to a significant interaction between the two factors (*F* _1,19_ = 10.90, *p* = .004, *η*^2^ = 0.026). Analyses using mixture modelling (Bays, Catalao, & Husain, 2009) confirmed that errors originating from responses to the non-cued grating orientation were rare (swap rate of .033 ± .01 on two-item trials; see also Huang, 2020).

**Table 1.** Descriptive statistics of error for all experimental conditions.

|  | Report 1st trials | Report 2nd trials |
| --- | --- | --- |
| 1 item | 11.91° ($\pm 4.61^\circ$) | 8.96° ($\pm 2.52^\circ$) |
| 2 items | 14.69° ($\pm 5.38^\circ$) | 9.79° ($\pm 2.89^\circ$) |

*Note.* Mean Absolute Error ( ± Std. Dev.). All units are in degrees.

### 3.2 Repulsive performance biases within trials

We analysed within-trial biases in behavioural performance by assessing whether the reported orientation was systematically reported as closer to or further away from the non-target orientation on the same trial (see **Methods**). We restricted the analyses to 2-item trials. **Figure 2A** shows the performance bias for all absolute angular distances between the first and second grating orientation for report 1st and report 2nd trials. In trials with report 1st cues, there was no significant bias towards or away from the interfering second grating orientation that was not relevant to the task at hand (*t*_19_ = 0.74, *p* = .467). In contrast, trials with report 2nd cues revealed significant biases away from the initially encoded first grating orientation (*t*_19_ = *−*2.33, *p* = .031; illustrated in **Figure 2B**). The repulsive bias in report 2nd trials was confirmed using a cluster-based permutation test, showing a significant cluster (p = .012) when the angular distance between the two orientations was between 10° and 49° (**Figure 2A**).

**Figure 2:**
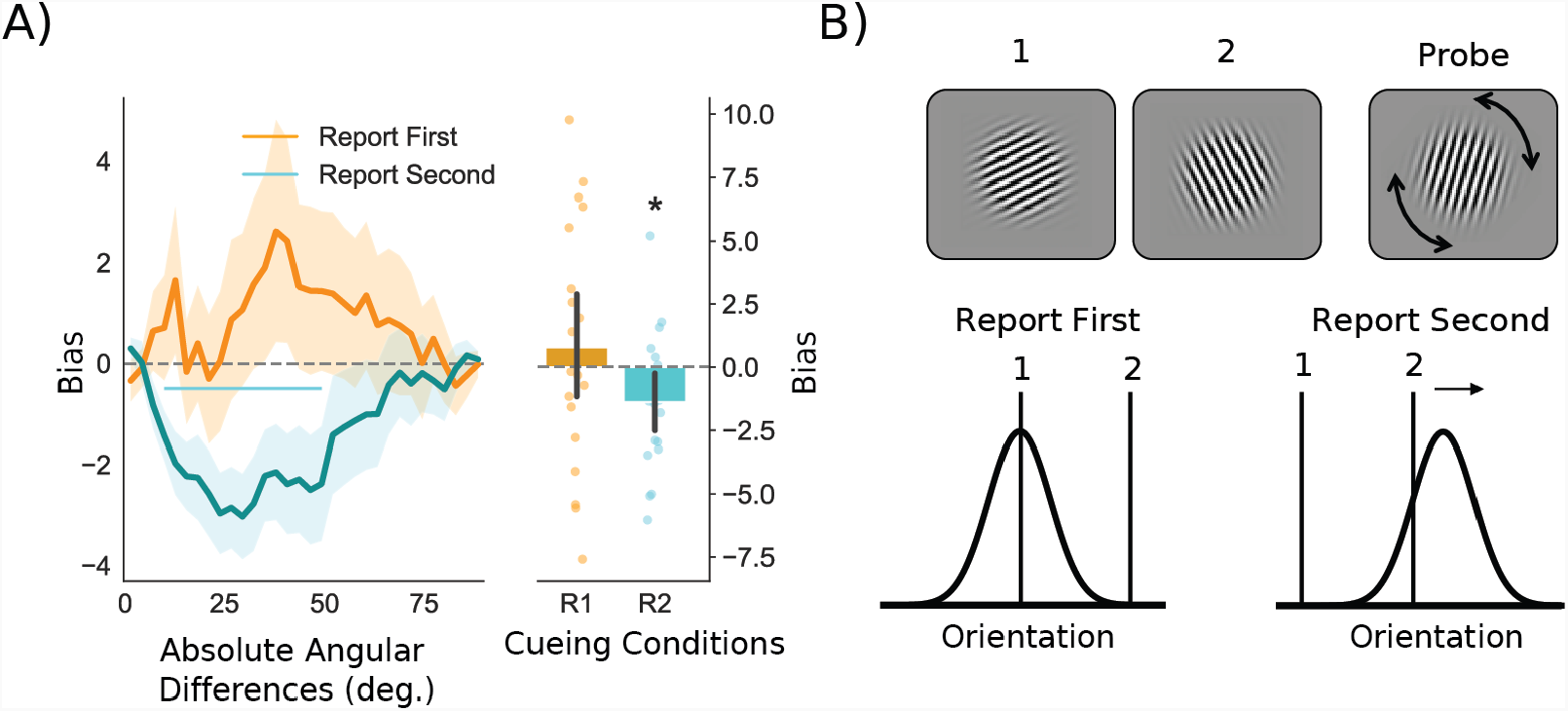
Within-trial repulsive performance biases. A) Within-trial bias as a function of the absolute angular distance between the two presented orientations. Trials were split for report 1st and report 2nd cues on 2-item trials. Shading indicates standard error of the mean. The horizontal line indicates a cluster of significant performance biases for a range of angular difference. The right panel shows the sum of bias across angular distances per subject. Error bars show 95% confidence intervals. R1 = report 1st; R2 = report 2nd. B) Schematic of the within-trial performance bias. A repulsive performance bias is observed for report 2nd trials only.

### 3.3 Attractive Performance Bias Between Trials

We next evaluated the between-trial bias on responses in the current trial towards the orientation that was cued on the previous trial (**Figure 3**). We assessed the performance bias as a function of angular differences between the target grating on the current and on the previous trial. The analysis also considered the position of the target grating in the current trial (1^st^ or 2^nd^) and the number of items on the current trial (1- or 2-items). For consistency, we term all trials where participants report the first grating orientation *report 1st trials* and trials where participants report the second orientation *report 2nd trials*, regardless of the number of gratings presented. Again, we calculated the sum of the bias across angular distances between targets in the current and previous trial **Figure 3A, B**. In contrast to the repulsive bias described in the previous section, we found that all conditions showed an attractive performance bias (all p < .05 in two-sided statistical tests). The attractive serial bias was most pronounced for small to intermediate angular distances between the inducer and current orientation (0 - 60°). A repeated-measures ANOVA on the sum of biases across angular distances indicated an effect of cue type, with larger biases occurring in report 1st trials (*F* _1,19_ = 5.706, *p* = .027, *η*^2^ = .172), but not of the number of gratings presented in a trial (*F* _1,19_ = .980, *p* = .335, *η*^2^ = .007). The two factors did not interact (*F* _1,19_ = .377, *p* = .547, *η*^2^ = .002). This shows that the bias was stronger when recalling the first item, which was encoded closer in time to the previous trial.

**Figure 3:**
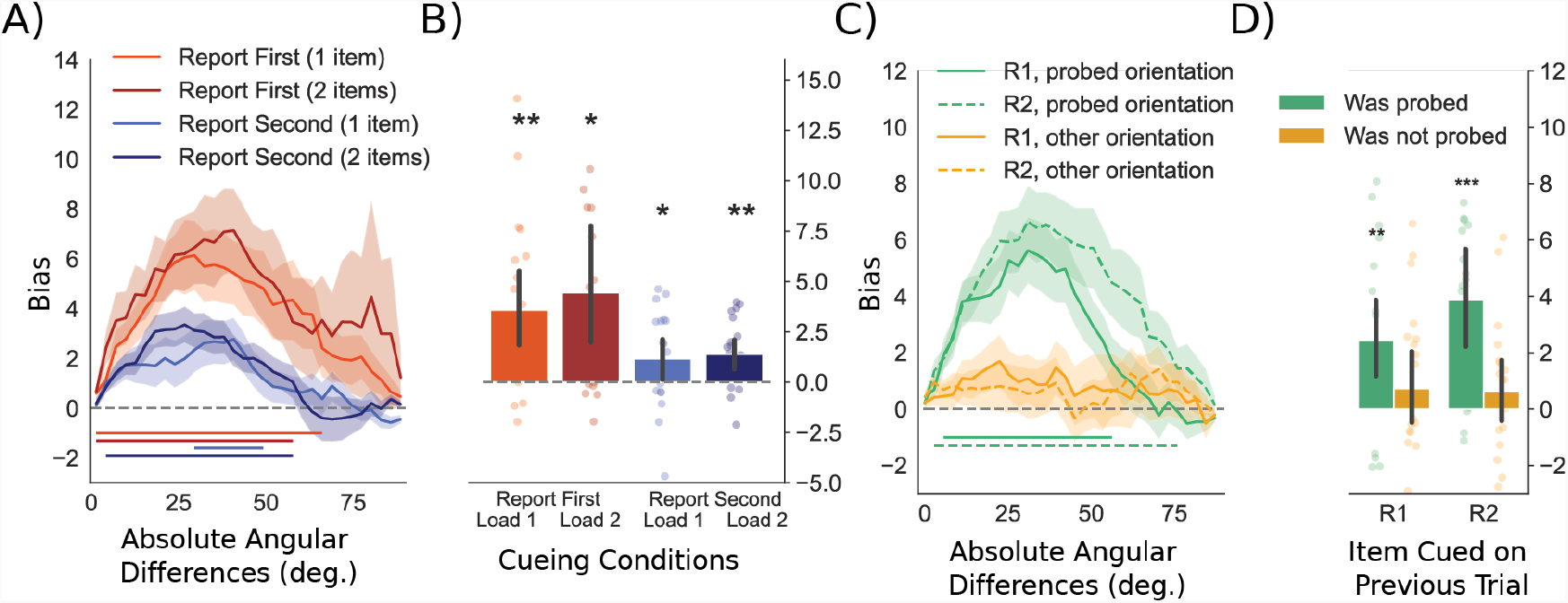
Attractive behavioural biases towards the relevant target orientation in the previous trial. A) Line plots for the attractive between-trial bias as a function of the absolute angular distance between the target orientation on the previous trial and the presented grating. Shading indicates the standard error of the mean. Horizontal lines indicate clusters of significant performance biases for a range of angular differences. B) Bar plots indicate the average sum of biases across angular distances per subject and per condition as seen in panel 3A. C) Line plots show the attractive between-trial bias as a function of the absolute angular distance between the probed angle on the previous trial (green) or the presented but not probed angle in the previous trial (orange). The horizontal lines indicate cluster-corrected angular distances for which the bias is significant. R1 = report 1st orientation on the previous trial; R2 = report 2nd orientation on the previous trial. D) Sum of the response data shown in panel 3C. Positive values indicate an attractive bias. All error bars indicate 95% confidence intervals.

### 3.4 Task Dependence of Attractive Performance Bias Between Trials

Next, we repeated the same between-trial analyses but investigated the role of task relevance and cue type in the previous trial. This allowed us to test for and compare behavioural biases elicited by the probed (and reported) orientation and by the unreported orientation in the previous trial. We also tested whether the cue type on the previous trial affected bias in the current trial. We only looked at trials in which two orientations were presented on the previous trial.

A repeated-measures ANOVA on the sum of angular distances of the performance bias for task relevance and cue type confirmed an effect of task relevance (*F* _1,19_ = 14.684, *p* =.001; **Figure 3D**) but showed no effect of cue type (*F* _1,19_ = 1.423, *p* = .248) or interaction (*F* _1,19_ = 1.633, *p* = .216). Task-relevant orientations in trials of either cue type resulted in a significant bias (previous trial was report 1st and contained 2 items: *t*_19_ = 3.524, *p* =.002; previous trial was report 2nd and contained 2-items: *t*_19_ = 4.476, *p <* .001). Unprobed orientations did not lead to a significant bias (both *p >* .2). No reliable difference was observed between the strength of the bias between report 2nd 2-items or report 1st 2-items conditions in previous trials (*t*_19_ = 1.691, *p* = .107). Following up, we assessed the performance bias as a function of the angular distance between the current target orientation and previously presented orientations. Cluster-based permutation testing showed an attractive bias towards the task-relevant orientation on the previous trial when a report 1st cue (*p* = .001; 7° – 58°; **Figure 3C**) or a report 2nd cue (*p <* .001; 4° – 77°) was presented in the previous trial. For unreported orientations, no bias was observed (no candidate clusters for report 1st or report 2nd cue). Together, this pattern of results is showing an attractive between-trial bias, but only with regard to items that were relevant in the previous trial.

### 3.5 Neural Classification of Presented Orientations

For our classification analysis, grating orientations were binned into 10 equally spaced bins. We applied Linear Discriminant Analysis (LDA) on spatial and temporal features from all 306 MEG sensors ranging from 400 ms prior to stimulus onset up to 900 ms post stimulus onset (see **Methods**). If orientation information was present, LDA likelihood estimations gave rise to representational similarity curves centred on the presented orientation that could be convolved with a cosine function to result in a single evidence estimation per time point. LDA classification reflected significant evidence for the presented orientation after the visual onset of the grating (**Figure 4**). This revealed significant decoding of both the first grating orientation (100 – 615 ms; *p <* .001) and the second grating orientation (95 – 590 ms; *p <* .001). A non-significant trend was observed for the evidence of the first grating orientation following the presentation of the second grating (250 – 600 ms; *t*_19_ = 1.971; *p* = .063; see also cross-decoding analyses below). There was no difference in classifier evidence after a report 1st or report 2nd cue (250 – 600 ms; *t*_19_ = 0.798; *p* = .435).

**Figure 4:**
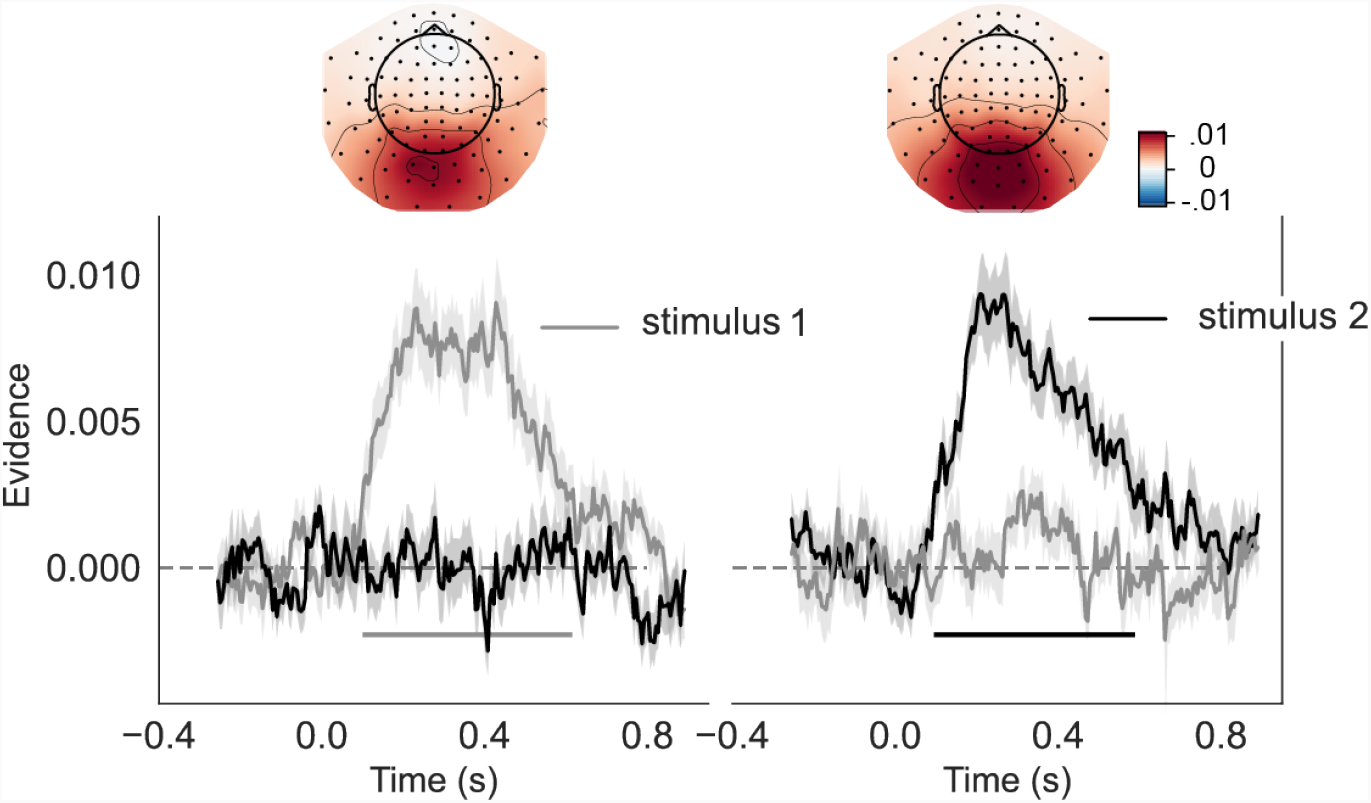
Orientation decoding of presented grating. Training an independent classifier to decode the first or second grating orientation in either the first- or second-time interval - left and right side respectively. The MEG topographies show which sensors most strongly contribute to the overall evidence. The left topography shows decoding of the first orientation in the first interval between 250 - 600 ms. The right topography shows decoding of the second orientation at the same latency. Error bars indicate the standard error of the mean. Horizontal lines indicate periods of significant grating orientation decoding after cluster-based permutation tests against zero, p = .05.

### 3.6 Within-trial Classification Bias Away From Previous Stimulus

Representational similarity curves were used to estimate the direction and magnitude of neural biases in orientation representation. By training the classifier on all stimuli, orientation history biases should cancel out, allowing testing on trials with specific prior orientations (clockwise vs. counterclockwise to current stimulus) to reveal any neural biases. To investigate how information from the second grating was modulated by information from the first grating, we separately assessed trials with report 1st and report 2nd cues. Doing so, we evaluated the classification evidence in the MEG data in the epoch following the presentation of the second grating. For these analyses, we again only selected trials with both gratings were presented. We separated trials in which the first grating was clockwise vs. counterclockwise relative to the second grating (angular distance of 10° - 50°, based on behavioural results, see **Figure 2A**, **B**). We considered the average of time points between 250 – 600 ms for all future analyses, since in this time window stimulus orientation could be decoded with reliable accuracy in this time window (see **Figure 4** and grey-shaded area in **Figure 5**). Echoing the performance biases, no significant bias occurred on report 1st trials (**Figure 5A, B**; *t*_19_ = *−*0.723; *p* = .478). However, on report 2nd trials, we observed a repulsive effect, away from the previously encoded grating orientation (**Figure 5C, D**; *t*_19_ = *−*3.52; *p* =.002).

**Figure 5:**
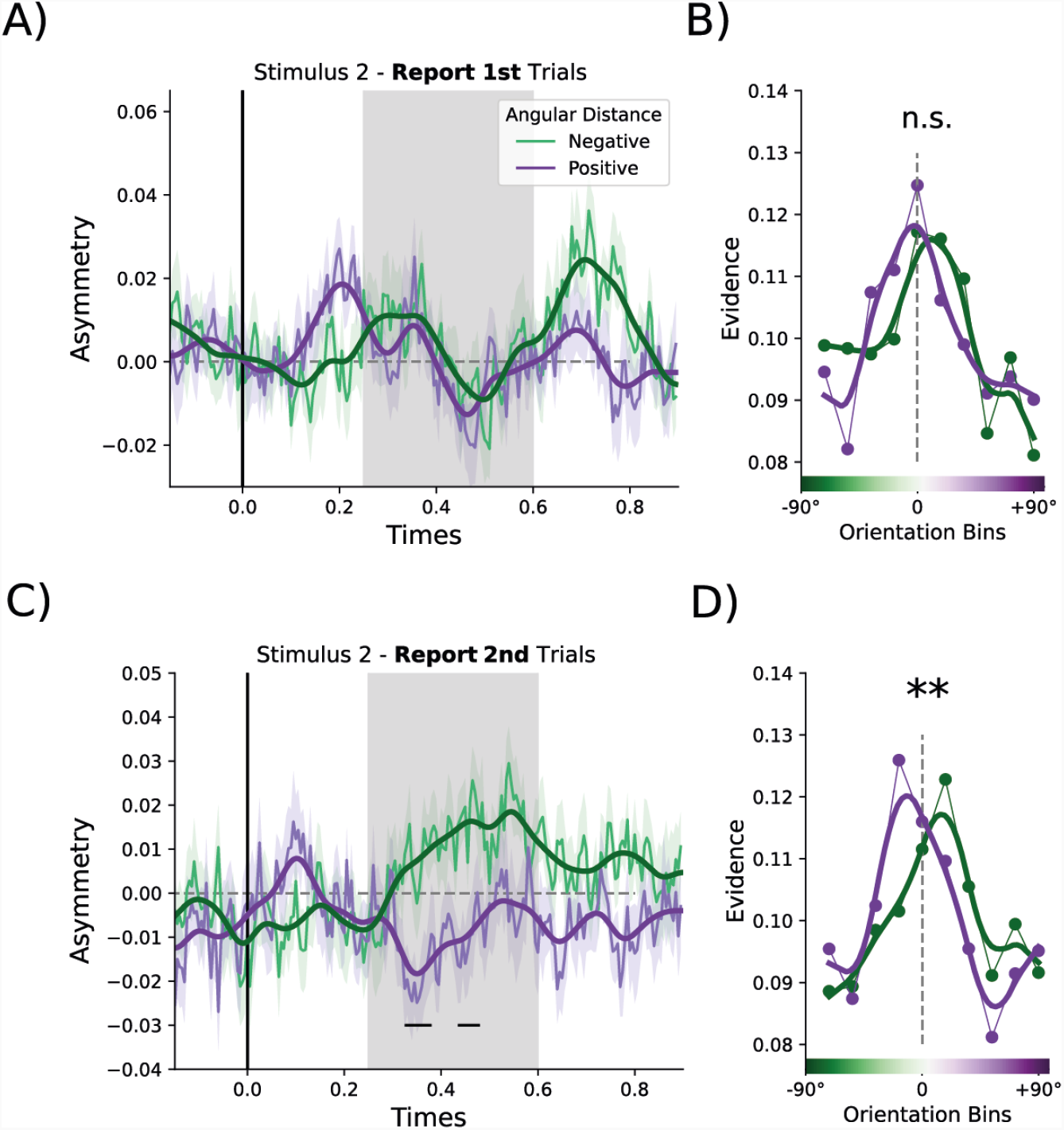
Shift in representational similarity curve relative to previous grating orientation. A) Asymmetry scores following the presentation of the second grating on report 1st 2-item trials. The first grating orientation had a positive (purple) or negative (green) angular distance relative to the current grating orientation. The shaded area indicates the time interval (250 - 600 ms) used for statistical analysis and to generate the representational similarity curves in the right panel. Smoothing (thicker lines in darker colours) was applied for visualisation purposes. B) shows the representational similarity curves with evidence for the presented grating orientation for trials with positive or negative angular distances relative to the first rating. For visualisation purposes, data points were interpolated (from 10 to 50 data points) and fitted using a Savitzky-Golay filter (with a window length of 9 data points and polynomial of order 1). C) Asymmetry scores for report 2nd 2-item trials, where participants encoded the grating orientation on screen into working memory with a neural bias away from the grating orientation that was presented earlier on the same trial. Shading indicates standard error of the mean. D) shows data from the grey-shaded region in panel 5C. Horizontal lines indicate periods of significant bias on unsmoothed data after cluster-based permutation tests against p = .05.

There was no correlation, across participants, between the magnitude of the bias in the behavioural and neural data on report 1st trials (*r* = .010; *p* = .968) or report 2nd trials (*r* = *−*.328; *p* = .158).

### 3.7 Repulsive Neural Biases Between Trials Away From Sensory History

To probe for neural between-trial biases, we used the same approach as for within-trial neural biases. We tested the LDA evidence derived during the stimulus encoding period for systematic deviations in likelihood estimations as a function of the angular distance between the current orientation and the probed orientation on the previous trial. If the behavioural between-trial bias reflects neural modulation during encoding of sensory features, we would expect to see an increase in likelihood for orientations presented in the previous trial, in line with the attractive performance bias. We trained the classifier on data following presentation of both the first and second grating orientation combined and included all cue types and number of items presented. Informed by our behavioural analyses on between-trial performance biases, for the test set we selected trials where the previous probe angle had a relative difference of 0° – 60° (**Figure 3A, B**; derived from significant angular differences) positive or negative from the presented grating orientation. The results were qualitatively the same and remained significant when other angular ranges were selected.

Contrary to our expectations, classifier evidence was significantly shifted away from the target orientation on the previous trial (250 – 600 ms post grating onset; *t*_19_ = *−*2.83, *p* =.011; **Figure 6A - D**) rather than mirroring the attractive behavioural bias. In practice, this would mean that if the cued orientation on the previous trial was CW, classifier evidence for CCW bins increased, and vice versa. The repulsive bias away from the target on the previous trial trended towards significance for the first (*t*_19_ = *−*1.78, *p* = .090) and was significant for the second grating (*t*_19_ = *−*2.43, *p* = .025) in the current trial when considered separately (**Figure 6C**). The between-trial repulsive bias during stimulus-two processing was present if no orientation was presented in the first interval (*t*_19_ = *−*2.69, *p* = .014) but not when the first grating was also presented (*t*_19_ = *−*1.68, *p* = .110). Interestingly, topographies in **Figure 6D** and **Figure 6G** were highly similar (*r* = .563; *p <* .001) and both topographies correlated negatively with the stimulus-decoding topography (**Figure 6D**, *r* = .558, *p <* .001; **Figure 6G**, *r* = .763, *p <* .001).

**Figure 6:**
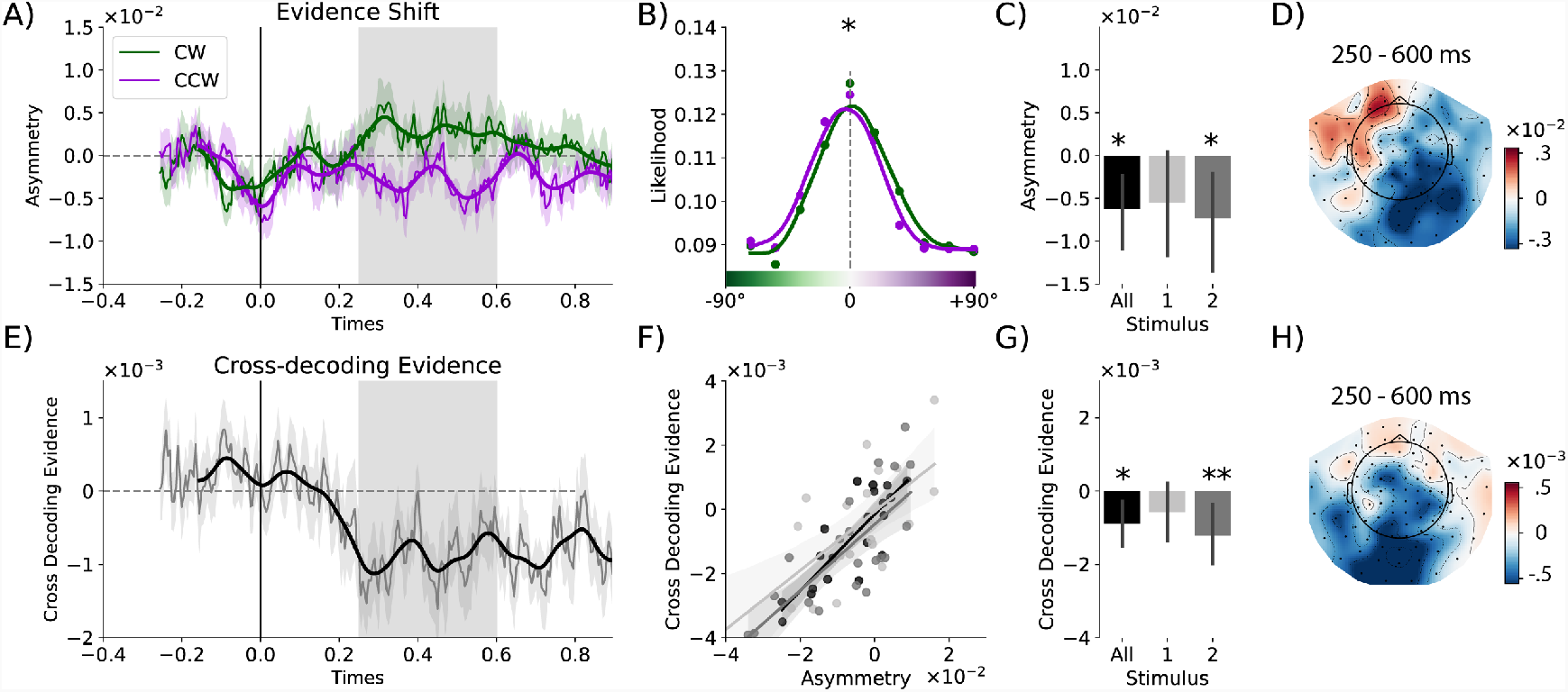
Bias on encoding imposed by previous trial. A) Asymmetry index for trials with a CW or CCW angular distance (0° – 60°) between the cued item on the previous trial and the presented grating (collapsed across the first and second grating). Asymmetry index time course is shown relative to the onset of grating presentation. A one-dimensional Gaussian filter with a kernel of 15 ms was applied to the data for visualisation purposes. The grey shaded area indicates which time points (250-600 ms) are used for analyses described in the text and in panels 6B-D and F-H. B) Average LDA orientation-likelihoods for trials with CW (green) and CCW (purple) angular distances with respect to the previous trial. Evidence is lower for CW orientation bins when in the previous trial a CW orientation was cued, and vice versa for CCW orientations. The same interpolation was applied as described in Figure 5. C) Average bias relative to the previously cued orientation across all stimuli and for the first and second orientation separately. D) MEG topography based on searchlight results, illustrating in which sensors the evidence shift from panels 6A-C is most prominent. E) Cross-decoding evidence for the previously cued orientation (classifier trained on current orientation), locked to grating presentation. Same conventions as in panel 6A. F) Correlations between asymmetry index panel 6A and cross-decoding evidence from panel 6E for grating orientation processing of all stimuli combined (black), first grating (light grey) and second grating (dark grey). A least-squares linear regression model was applied for each of the conditions, with shaded 95% confidence intervals estimated using bootstrapping. G) Cross-decoding evidence from panel 6D summarised for all stimuli and separately for the first and second grating. H) MEG topography based on searchlight results, showing cross-decoding evidence for each sensor. *: p < .05, **: p < .01. Plotted with 95% confidence intervals.

Next, we tested whether this repulsive neural bias was affected by the task relevance of the grating and by the cueing condition of the previous trial. We quantified this bias for task-relevant and task-irrelevant grating orientations in the previous trial. Only trials where the previous trial contained two items were included in this analysis. When averaging the neural bias over 250 – 600 ms, the task-relevant orientation showed a repulsive neural bias (t_19_ = -2.78, p = .012), but we found no neural bias for task-irrelevant orientations (*t*_19_ = *−*0.33, *p* = .743), though this difference did not reach significance (*t*_19_ = 1.80, *p* = .088). Yet, cluster-based permutation testing indicated a significant cluster indicating that task-relevant orientations exerted a significantly stronger repulsive bias than task-irrelevant orientations (500 – 550 ms; *p* = .039).

### 3.8 Cross-Decoding Evidence for Previously Presented Stimuli

If information about the previously presented orientation is still partially present in the visual system during and after the presentation of the current grating orientation, it could interact with encoding, possibly leading to the observed repulsive bias. One possibility is that the lingering representation is in an orthogonal representational format, which is different from sensory coding of features (Libby & Buschman, 2021). In this case there would be little to no overlap between the activation pattern elicited during sensory input and the pattern related to the lingering representation of that past grating orientation. A classifier trained to separate perceptual information would therefore not cross-generalise if tested on the memory code. Alternatively, the lingering code could be present in a stable representation that shares similarity with the representation of incoming sensory information. If this were the case, a classifier, trained on the data from the current grating orientation, would cross-generalise and identify information about the past grating (or suppression of information expressed in negative evidence).

We adapted cross-decoding to detect lingering orientation-selective activity from the previous trial (also see Wan, Cai, Samaha, & Postle, 2020). After training the classifier for the presented orientation and testing for evidence of the previous trial’s target orientation, we observed significant negative classifier evidence in the period of 250 – 600 after grating onset (**Figure 6F**; concatenating over grating one and grating two: *t*_19_ = *−*2.820, *p* =.011; grating one alone, *t*_19_ = *−*1.376, *p* = .185; grating two, *t*_19_ = *−*2.951, *p* = .008).

Negative classifier evidence indicates that, while information is still present about the previous orientation, orientation-selective patterns may be sign-reversed relative to stimulus encoding. This suppression of evidence for the previous trial’s orientation could have been the cause of the apparent repulsive bias in the decoding of the current trial’s orientation. If this was the case, we would expect the two measures to be positively correlated: stronger suppression of the previous orientation (negative classifier evidence) should lead to a more negative bias for the current orientation. We tested this using Pearson correlations across participants and found a significant correlation when assessing all grating presentations together (**Figure 6E**; *r* = .832, *p <* .001; grating one alone, *r* =.685, *p <* .001; grating two alone, *r* = .751, *p <* .001).

No correlation was observed between the attractive behavioural performance bias and the magnitude of the negative shift in the neural data (*r* = .186, *p* = .432) nor with the magnitude of negative decoding (*r* = .144, *p* = .544).

## 4 Discussion

The present study investigated biases from previously perceived and memorised information on neural coding and behavioural responses. We observed both neural and behavioural biases that demonstrate interactions between past and present sensory processing. Orientations presented on the *same* trial exerted repulsive performance biases on currently perceived orientations. In contrast, task-relevant orientations on the *previous* trial exerted an opposite, attractive performance bias, altogether providing evidence for two counteracting biasing processes acting in tandem. Interestingly, multivariate decoding of neural data indicated that both stimuli from the same and previous trial generate a repulsive neural bias, suggesting that the two types of performance biases may arise from modulations acting on different stages of stimulus processing. While repulsive biases could reflect mechanisms akin to visual adaptation that promote visual discriminability, the attractive performance bias observed across trials had no equivalent attractive modulation at the level of the sensory representation. Since a neural repulsive bias occurred instead, we suggest that post-perceptual modulatory mechanisms may override any early repulsive sensory modulation and lead to attractive performance biases.

The present behavioural results provide evidence of two types of performance biases: a repulsive within-trial and an attractive between-trial performance bias (c.f., Bae & Luck, 2020; Fischer, Czoschke, et al., 2020). Both biases were modulated by the task relevance of the inducing stimulus. Firstly, we identified a repulsive performance bias away from the first orientation in a trial, but no significant retrospective repulsive bias from the second orientation, even though the second grating was presented closer in time to the probe. We speculate that this may have happened because the relevance cue appeared together with this second stimulus, indicating its irrelevance in probe-first trials. Similarly, participants’ responses in the current trial were biased only towards task-relevant orientations in the previous trial. At the neural level, biases were also dependent on task relevance. A repulsive neural bias was only present when the presented orientation was cued as task-relevant and therefore encoded into working memory. By contrast, task-irrelevant gratings that were not encoded into working memory could be decoded with similar precision but did not exhibit a significant bias. Gratings from previous trials that were associated with an attractive performance bias also led to repulsive neural biases during working-memory encoding. Only task-relevant orientations led to a repulsive bias.

Adaptation could partly explain the repulsive neural bias observed here. Visual adaptation has been proposed as the cause of repulsive neural biases (Jazayeri & Movshon, 2006, 2007; Kohn, 2007; Stocker & Simoncelli, 2008; Webster, 2015). Since adaptation reduces firing in recently active neurons (Clifford, Wenderoth, & Spehar, 2000; Wainwright, 1999), it is an efficient use of finite neural resources (Stocker & Simoncelli, 2008; Webster, 2015) when the environment is autocorrelated because neurons can code for a larger range of stimuli when their responses are not saturated. Curiously, we observed a repulsive neural bias only on report 2nd trials, when the presented orientation was task-relevant and encoded into working memory. The interaction with task relevance suggests that stimulus processing is only biased when it is primed for later use in upcoming behaviour.

This task-dependent modulation was unlikely the result of reduced processing of task-irrelevant stimuli, as overall orientation decoding was not affected by task relevance. In line with recent studies, it is possible that a context-sensitive repulsive bias, possibly occurring at a post-perceptual stage (Fritsche & de Lange, 2019; Zamboni, Ledgeway, McGraw, & Schluppeck, 2016), exists alongside an early perceptual bias based on visual adaptation (Fritsche, Mostert, & de Lange, 2017). The task-dependency of the neural bias could be the basis of recently observed repulsive biases in behaviour (Bae & Luck, 2017; Chunharas, Rademaker, Brady, & Serences, 2019; Czoschke, Fischer, Beitner, Kaiser, & Bledowski, 2019; Czoschke, Peters, Rahm, Kaiser, & Bledowski, 2020), which may help individuate concurrently maintained stimuli (e.g. Wei, Wang, & Wang, 2012). Under this explanation, only attended and encoded features would be subject to interactions with previous features.

The bias imposed by the orientation from the previous trial was attractive in behaviour but repulsive in the neural data at early processing stages. This contrast is ostensibly at odds with previous behavioural studies that have assigned an early perceptual origin to attractive between-trial biases (e.g., Cicchini, Mikellidou, & Burr, 2017; Cicchini, Mikellidou, & Burr, 2018; Fischer & Whitney, 2014). There is still little direct neural evidence confirming this. EEG studies have shown that previous trial information can be decoded during the encoding phase of the current trial (Bae & Luck, 2019) or immediately prior to the current trial (Barbosa et al., 2020), and visually evoked neural responses in numerosity judgement tasks are modulated by stimulus history (Fornaciai & Park, 2018, 2019). However, the mere presence of prior stimulus information does not imply an attractive bias on the current stimulus. One study observed an attractive behavioural bias and a neural bias in early visual areas using fMRI (St John-Saaltink, Kok, Lau, & de Lange, 2016). However, this study used only two stimulus orientations (45° and 135°) with an offset too large to produce a reliable behavioural bias, meaning that our findings may reflect a different biasing phenomenon.

Contradicting the early sensory origin of serial attractive bias, a recent fMRI study (Sheehan & Serences, 2021) found evidence consistent with the present results. Sheehan et al. observed repulsive neural biases relative to the orientation on the previous trial across in visual cortex, in spite of an attractive behavioural bias, and found that models incorporating early visual adaptation and a post-perceptual origin of attractive biases could explain both effects. These results are in line with the present findings. Together, the observations make a case that prior stimuli lead to repulsion at the encoding stage and that attractive performance biases may arise elsewhere. Our findings further indicate that the link between neural adaptation and behaviour should be context-dependent, since we observed repulsive neural biases together with both repulsive *and* attractive behavioural biases. Therefore, future models linking visual adaptation to behaviour may need to incorporate context-dependence. Additionally, the high temporal resolution of MEG allowed us to show that neural biases arise within 500 ms of stimulus onset, have a posterior origin, and that they occur simultaneously relative to multiple prior stimuli (from the same trial and the previous trial).

While Sheehan and colleagues argued that past stimuli were stored in a non-sensory code, they did not directly examine whether the representation of past stimuli occurred in a shared neural subspace with the representation of current stimuli. Here, we addressed this issue by showing that neural suppression of recently active neural populations could account for the observed repulsive bias. We tested this using cross-decoding analyses, training a classifier on the presented orientation and predicting previous orientations. Consistent with the suppression of recent stimulus-specific activity, cross-decoding yielded below-chance decoding of the previous orientation. In turn, this may have shifted the neural tuning curve for the current orientation away from the previous orientation, generating a repulsive bias. This relationship was confirmed by the robust correlation between cross-decoding and repulsive bias magnitude.

Another recent study observed a repulsive neural bias in single-unit recordings from frontal eye fields (FEF) paired with an attractive behavioural bias in a delayed-saccade task (Papadimitriou, White, & Snyder, 2017). The authors suggested that lingering attention to the previous target location could warp the representation of current target locations (Zirnsak, Steinmetz, Noudoost, Xu, & Moore, 2014). Since we observed neural biases primarily in posterior sensors, our results are more in line with a sensory origin of the repulsive bias, but this may interact with attentional biases originating in frontal cortex (Moore & Armstrong, 2003; Taylor, Nobre, & Rushworth, 2007). Attentional modulation could be one explanation for the context-dependency of biases observed here.

Altogether, we demonstrate a consistent repulsive shift in neural evidence during working-memory encoding. Our results imply that perceptual adaptation, along with context-sensitive factors, contributes to feature-selective downweighting to exert a repulsive bias away from recent stimulus features. Interestingly, no evidence of an attractive neural bias acting directly on sensory aspects of encoding was observed. Neural data thereby provide indirect evidence for the post-perceptual account of attractive between-trial biases, rather than modulating encoding stages (Bae & Luck, 2020; Bliss, Sun, & D’Esposito, 2017; Fritsche, Mostert, & de Lange, 2017; Kim, Burr, Cicchini, & Alais, 2020; Pascucci et al., 2019). We speculate that the attractive between-trial bias instead arises through post-perceptual processing stages involving memory (Bliss, Sun, & D’Esposito, 2017; Fritsche, Mostert, & de Lange, 2017), perceptual decision-making, or motor planning (Boettcher, Gresch, Nobre, & van Ede, 2021; de Azevedo Neto & Bartels, 2021; Sadil, Cowell, & Huber, 2021). The source of the attractive between-trial bias, whatever its neural mechanism, may be strong enough to override the repulsive bias during the perceptual/encoding stage. Together, these co-existing biases may help guide efficient coding for nuanced perceptual discriminations and visual stability across our environment.

## Conflict of interest

The authors declare no conflict of interest.

## Acknowledgements

This research was funded by an ESRC Grand Union studentship and the Scatcherd European Scholarship awarded to **J.E.H**., an ERC Starting Grant from the European Research Council (MEMTICIPATION, 850636) to **F.v.E**., was supported by a James S. McDonnell Foundation Scholar Award (220020405) and an ESRC grant (ES/S015477/1) to **M.G.S**., a James S. McDonnell Foundation Understanding Human Cognition Collaborative Award (number 220020448) and a Wellcome Trust Senior Investigator Award (104571/Z/14/Z) to **A.C.N**., as well as a Wellcome Trust award (201409/ Z/16/Z) and with support from University College Oxford to **N.E.M**. The work was enabled by the NIHR Oxford Health Biomedical Research Centre and the Wellcome Centre for Integrative Neuroimaging is supported by core funding from the Wellcome Trust (203139/Z/16/Z). The funders had no role in study design, data collection and analysis, decision to publish, or preparation of the manuscript. For the purpose of Open Access, the author has applied a CC BY public copyright licence to any Author Accepted Manuscript version arising from this submission.

## Data Availability

Data and analysis scripts can be accessed via https://osf.io/98ujm/.

## Notes

### Competing Interest Statement

The authors have declared no competing interest.

### Summary of Updates

Figures updated, Introduction and Discussion updated, terminology made more consistent.

